# Pace of life predicts parasite resistance and fecundity tolerance, but not mortality tolerance, among Trinidadian guppies, *Poecilia reticulata*

**DOI:** 10.64898/2025.12.17.694895

**Authors:** Sadie Evanov, Faith Rovenolt, Natalia Tepox-Vivar, Rina Zambetti, Kelley Troutman, Vineet Nayak, Kaylee Mulligan, Saisreetha Midde, Noah T. Leith, Jessica F. Stephenson

## Abstract

Host defence against parasites can include limiting parasite growth, ‘resistance’, limiting the mortality cost of infection, ‘mortality tolerance’, and limiting the reproductive cost of infection, ‘fecundity tolerance’. Theoretically, these three host strategies have very different epidemiological and evolutionary outcomes. In particular, because mortality tolerance increases parasite population size, it is under strong positive frequency-dependent selection and may therefore be less variable between populations than either resistance or fecundity tolerance. Additionally, host investment in each strategy can be expected to differ between populations that experience different ecological conditions. Here, we tested how populations of Trinidadian guppies *Poecilia reticulata* from the upper and lower courses of three rivers responded to experimental infection with a novel strain of *Gyrodactylus turnbulli*. In line with theoretical predictions, we found that lower course populations, previously shown to have faster paces of life, invested less in resistance and fecundity tolerance - but not mortality tolerance - than the upper course populations with slower paces of life. Our results indicate that this host-parasite interaction both conforms to evolutionary-epidemiological theoretical predictions, and is shaped by broader ecological conditions.

## Introduction

In this epoch of infectious disease emergence and ecological change, understanding the evolution of host defences, particularly across ecological contexts, is essential [1]. Host defence against parasites can take several forms, each with its own implications for host and parasite fitness, and thus evolutionary and co-evolutionary dynamics [2]. Such defences are typically categorised as either ‘resistance’, if hosts limit parasite growth, or ‘tolerance’, if hosts limit the fitness cost of infection without affecting parasite growth. Intuitively, these two broad categories of defence should have different epidemiological and evolutionary implications [3–5]. Specifically, as resistance reduces parasite fitness and thus disease prevalence, selection on resistance is negatively frequency dependent. The implications of tolerance depend on the type: ‘fecundity tolerance’, in which hosts limit the reproductive costs of infection, does not necessarily impact disease prevalence. By contrast, ‘mortality tolerance’, in which hosts limit the survival costs of infection, may increase disease prevalence as infected hosts live - and can therefore transmit for - longer, leading to positive frequency dependence. Epidemiological feedbacks may therefore promote diversity in resistance, not affect diversity in fecundity tolerance, and limit diversity in mortality tolerance. These predictions have been broadly supported by theoretical [6–8] and empirical work across systems [2,9,10].

In parallel, the broader community with which both host and parasite may interact can importantly affect the evolutionary trajectory of host defences. In particular, predation pressure may constrain or redistribute host investment in defense against parasites through at least two pathways. First, high predation pressure can favor individuals that maximize resource allocation into fecundity and reproduction early in life (i.e., ‘fast’ pace of life) [11,12], at the expense of reduced investment in both resistance [13,14] and tolerance [15,16] of parasites, in line with predictions [17,18]. Populations with fast life histories may thus show weaker fecundity tolerance even though they have evolved increased fecundity overall. Second, high-predation environments select for antipredator behaviours like social grouping, which may increase the opportunity for parasite transmission, leading to increased parasite population size [19] and therefore stronger selection for investment in host defences. Thus, the outcome of selection on host defences due to epidemiological feedbacks likely depends on the ecological context hosts experience - of which predation pressure is a key component.

Here, we used females of the guppy, *Poecilia reticulata*, from Trinidadian populations under different predation regimes, and its gyrodactylid parasite *Gyrodactylus turnbulli*, to explore patterns of diversity in parasite resistance, fecundity tolerance, and mortality tolerance. Due to waterfalls that block the upstream migration of aquatic predators, guppies from the upper courses of Trinidadian rivers typically experience lower predation pressure than the lower courses [11,20]. This difference has resulted in evolved divergent life history strategies between upper and lower course guppy populations: lower course guppies tend to have faster paces of life than upper course guppies [11,20]. Field surveys show that lower course guppies are more likely to be infected, with more individual *Gyrodactylus* spp. [21], and that infected fish are in worse condition than uninfected fish in lower but not upper course populations [15], consistent with slower pace of life populations investing more in both resistance and tolerance. However, these studies leave unclear whether their findings reflect population-level differences in host defence, or the fact that lower course *Gyrodactylus* spp. are more virulent [19]. Therefore, to understand population-level variation in host defences, and how this might follow theoretical predictions across ecological contexts, we present a common-garden, holistic comparison of host defences across Trinidadian guppy populations.

## Methods

### Fish and parasite origin and maintenance

For this study we used female guppies only: as guppies are internally fertilising livebearers, fecundity tolerance is difficult to measure among males in this system. Females were laboratory-reared descendants of guppies collected from the upper (n = 19; 3 rivers) and lower (n = 27; 2 rivers) courses of the Guanapo, Aripo, and Yarra rivers in Trinidad in March 2020. Lower Yarra was not available. Stocks were housed and bred in mixed sex 75 L tanks on a recirculating system (Aquaneering) with daily 20% water exchanges, 12-hour L:D light cycles. The fish were fed daily with Tetramin flake and *Artemia*. Water hardness, pH, conductivity, and temperature were monitored daily, and kept at 117-135 ppm, 7.2-7.8 pH, 550-750 μS, and 25□±□1°C respectively. Prior to the experiment, females were removed from stock tanks and housed individually in 1.8L tanks under the same conditions.

To establish our parasite line, we transferred a single *G. turnbulli* individual from a commercially obtained guppy to a mixed stock, naïve, laboratory-bred host. We maintained this line on a group of naive hosts - ‘culture fish’ - in a 1.8L tank under standard conditions as above. Culture fish were screened twice a week under a dissection microscope, and naïve fish were added as needed to maintain the parasite population.

### Experimental infections

Experimental infections were conducted between September 12th - November 13th, 2023, in three batches. On day 0 of each batch, each experimental fish was anesthetized using tricaine methanesulfonate (‘MS222’; 4□g/L) in a crystallising dish. Once immobile, fish were oriented laterally with their left side facing upwards and a photograph including a ruler was taken using a mobile phone camera. These images were subsequently used to measure the fish standard length to the nearest 0.1 mm in ImageJ [22]. The fish was then poured onto a paper towel, gently dried on both sides, and weighed to the nearest 0.001 g. Next, the fish was placed back in a crystallizing dish in close contact with a recently euthanized, highly infected donor culture fish until approximately 2 worms transmitted (mean±SEM: 1.97±0.14), observed under a dissecting microscope. After infection, fish recovered from anesthesia in 1.8 L tanks in which they were kept individually under standard conditions for the remainder of the experiment. Wastewater passed through multiple filtration systems before being distributed to other tanks, preventing parasite transmission between them.

We quantified parasite loads every Monday, Wednesday, and Friday from day 1 to day 17. Fish were anesthetized for no longer than a total of five minutes for each count. Under a dissecting microscope, we manipulated each fish using a micropipette tip to accurately count the parasites on all body parts. After counts, fish were moved to fresh water and returned to their isolation tanks once they had recovered from anesthesia. Fish that died before the end of the experiment were individually preserved in 70% ethanol. After counts on day 17, fish were weighed and measured as described above, and euthanized using an overdose of MS222 before being preserved in 70% ethanol.

### Dissections

To quantify the number of offspring the experimental females were carrying, we dissected them in crystallizing dishes under a microscope. Using dissection scissors, we made a small incision behind the gravid spot on the ventral side of the female and extended to the bottom of the operculum. Two shallow vertical cuts were made on either end of the incision on the dorsal side of the female to create an opening into the abdominal cavity of the fish and expose the uterus, as is standard in the literature [23]. Any embryos were removed from the uterus and counted.

### Data Analysis

All analyses were conducted using R statistical software (v 4.4.0) [24]. We used *gpairs* [25] to visualise the data, *DHARMa* [26] to assess model fit, *car* [27] and *emmeans* [28] to extract statistics, and *visreg* [29] and *ggplot2* [30] to plot the figures. The supplementary material includes the complete code and output.

We used the data to address three questions. For the first, ‘do the courses differ in resistance?’, we used a linear mixed effects model (LMM) with the area under the curve of infection load over time up until day 11 of infection (‘infection severity’, using *MESS* package [31]) as the log-transformed response variable. As fixed effects, we included body length, number of offspring, river course, and how long the fish lived during infection (‘infdeathdays’; to control for death-induced inaccuracy in the infection severity variable: parasite growth and our counts cease once the fish has died). As random effects, we included the identity of the donor used to infect the fish, its river of origin, and its population of origin.

We addressed our second question, ‘do the courses differ in fecundity tolerance?’, using a generalised linear mixed model (GLMM) with a Poisson error distribution and log link function. We used the number of offspring a female was carrying as the response variable, and included as fixed effects body length, river course of origin, infdeathdays, infection severity, and - to test for differences in tolerance between the courses - the interaction between infection severity and river course. The random effects were as described above.

We addressed our third question, ‘do the courses differ in mortality tolerance?’, using a GLMM with a binomial error distribution and logit link function. We used whether or not the fish survived until the end of the 17-day infection period as the response, and included as fixed effects number of offspring, body length, course, infdeathdays, infection severity, and again the interaction between infection severity and river course. As random effects, we included the river and population of origin (donor identity caused model fitting issues). We also addressed this question using a mixed effects cox model looking at survival over the course of the 17 day infection, with the same fixed and random predictors as the GLMM.

## Results

Our ‘resistance’ model revealed that fish descended from upper and lower course populations differed in the average severity of their infections (our proxy for resistance; Fig. 1A; □ ^2^=4.33, df=1, P=0.037). As is common in this system, larger fish had more severe infections (□ ^2^=6.55, df=1, P=0.010), as did those that lived longer (because of the nature of the data collection, □ =8.46, df=1, P=0.004). We also found that donor identity explained ∼35%, and river of origin explained ∼30% of the variance in infection severity.

**Fig. 1:**
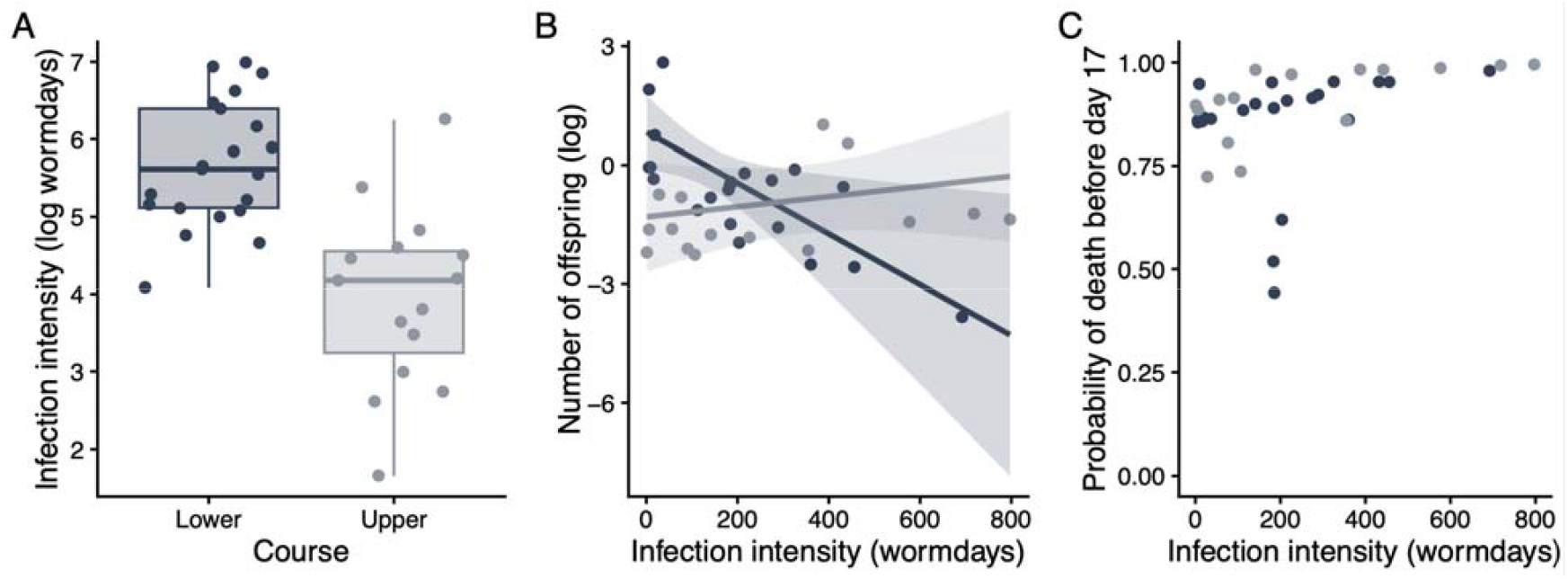
Guppy populations differ in parasite resistance (A) and fecundity tolerance (B), but not mortality tolerance (C). The points on all panels are the partial residuals from models described in the text. Panel A gives the median (thick line), interquartile range (box), and values within 1.5 x the interquartile range (whiskers). Panel B gives the model fit (lines) and 95% confidence interval (bands). There was no significant pattern in the data presented on panel C.

Our ‘fecundity tolerance’ model showed that fish descended from upper and lower course populations differed in the relationship between infection severity and number of offspring, consistent with upper course fish being more tolerant of infection (Fig. 1B; infection severity:course interaction: □ ^2^=5.16, df=1, P=0.023). Post-hoc tests confirm that the slope is significantly negative among lower course fish (*z*=-2.320, P=0.02), and not different from 0 among upper course fish (*z*=0.802, P=0.422). Fish that lived longer also tended to be carrying more offspring (□ ^2^=7.14, df=1, P=0.008). The random effects explained negligible amounts of the variance.

Our ‘mortality tolerance’ GLMM and survival analyses both showed that there was no difference between fish descended from upper and lower course populations in how infection severity affected their probability of dying before the end of the experiment (Fig. 1C; infection severity:course interaction: GLMM □ ^2^=0.075, df=1, P=0.785; Cox □ ^2^=0.820, df=1, P=0.365). Both models found that smaller fish were more likely to die (GLMM □ ^2^=0.075, df=1, P=0.785; Cox □ ^2^=0.820, df=1, P=0.365), and no other predictors explained significant portions of the variation. Full model outputs and validation are provided in the supplementary material.

## Discussion

Consistent with theoretical work and empirical work from other systems, our results support the maintenance of variation across populations in both parasite resistance and fecundity tolerance, but not mortality tolerance. We additionally show that investment in resistance and fecundity tolerance follow the pattern predicted by variation in predation pressure and pace of life theory across natural populations. We necessarily excluded males in this study which, because the sexes interact with these parasites differently across both ecological [15,21,32] and evolutionary [33,34] processes, may limit our ability to extrapolate from these findings. However, that the results from this laboratory experiment concur with data from field surveys suggests that the patterns we observed here are relevant to the system in nature.

We found that upper course guppies are more resistant than lower course guppies, which is in line with some previous observations in this system [21,35] but not others ([36,37]; using Aripo guppies, as we did here). Consistent with this variation, we found that river of origin and the identity of the donor explained substantial portions of the variance, emphasising that maximising sample size across these levels was important to uncover the general relationship we show here. Aspects of the donor’s identity have affected parasite growth on experimental fish in a previous study ([38]; in this case, whether the donor had transmitted before), but the mechanisms remain unclear. Overall, this result is consistent with investment in resistance that follows population differences in predator-driven life-history evolution [15,17,18].

We did not find support for the alternative hypothesis that predator-induced increased exposure to parasites selecting for increased investment in apparently costly parasite resistance, possibly due to a ‘resistance is futile’ effect [39], in these heavily parasitised lower course populations. That we found variation between populations in resistance suggests that it has costs which, combined with negative frequency dependent selection, prevents fixation [6,40,41]. Indeed, when mated to parasite naive females, male guppies that experienced lower infection severities produce offspring slower than those that experience heavier infections [32]. Such a cost would be especially detrimental to lower course guppies that experience higher extrinsic mortality from both predators and parasites [19].

While the patterns we observe are in line with evolutionary predictions, the extent to which the variation we found is genetic is unclear: all three defence metrics likely have some degree of plasticity. In other systems, fecundity responds to resource availability [42] and to infection in nuanced ways: infection can result in lower [32,43], higher [44], or ecomorph-dependent changes [45] in reproductive output. Similarly, host immune investment responds plastically to exposure to predators [46]. However, while we observe similar plasticity in this system [32,47], we know that both resistance [32,34,35,48] and tolerance ([34]; quantified using body condition as a proxy for fitness) are heritable, and we kept experimental conditions as consistent as possible.

We present our fecundity tolerance result as infection affecting fecundity, but we cannot rule out the effects of pregnancy on immune function. Indeed, pregnancy represents a life-history state in which the costs of immune activation increase [4,49]. A potential alternative explanation of our fecundity tolerance result is therefore that females vary in their ability to acquire resources: some lower course guppies appear able to invest in both reproduction and immunity, keeping parasite loads low while producing many offspring [50]. We did find that females carrying more offspring lived longer, potentially further corroborating the idea that some females were overall higher quality, and able to carry larger broods while both resisting and tolerating infection.

However, the population-level differences we observed under common garden conditions are not consistent with this explanation.

Guppy biology is such that we could not know what stage of their reproductive cycle females were in when we infected them. Female guppies are lecithotrophic: embryos are nourished by yolk before birth, with no additional maternal provisioning during development [51]. Instead, female guppies invest shortly before parturition: a new batch of ova matures and is fertilized after a brood is born [51]. This investment is substantial: reproducing females allocate approximately twice as much energy to reproductive tissues at the expense of somatic growth compared to virgins [52]. In our experiment, females were taken from mixed-sex breeding tanks, and all were sexually mature adults. It is highly likely that they had all mated with at least one male before being isolated and infected. However, as females store sperm [20], and exhibit cryptic female choice [53] they likely control when and how many eggs are fertilised, and potentially embryo developmental speed [54]. It therefore almost impossible to know at which stage in their reproductive cycle they were when infected, and likely would be even had we employed a controlled mating design.

Dissecting the females increased the accuracy of our fecundity metric, but it remains imperfect. Females under stress can prematurely birth offspring [55], and newborn fish are vulnerable to cannibalism, particularly in small tanks such as those we used [56]. Despite females being isolated and monitored daily for at least 20 days, we only noted one birth during the experiment (removing this female did not change our results), potentially indicating that additional births were missed due to cannibalism. Further, while there is no evidence for selective abortion in guppies or even matrotrophic poecilids under starvation [57,58], abortion and resorption under the stress of infection in this experiment may have allowed females to divert resources towards immunity.

Overall, our results support the role of epidemiological feedback in the evolution of host defences, and highlight that host defence evolution is additionally subject to selection from interactions with the broader ecological community.

## Data accessibility

The data are archived on Dryad, and can be accessed by reviewers here http://datadryad.org/share/LINK_NOT_FOR_PUBLICATION/a1KjYcZBR0w3-8fw9m7aqmWm9oYYIw1ypmRSKAzdyvQ.

## Ethics statement

This work was approved by the University of Pittsburgh’s Institutional Animal Care and Use Committee (IACUC), protocol 24065067.

## Author contributions

SE, RZ, FR and JFS conceptualised this study; FR, KM, KT, VN, SM and NTV collected the data; SE, RZ, NTL and JFS analysed the data; JFS, FR, SE wrote the first draft; all authors revised the manuscript and approved its submission.

## Acknowledgements

We thank Carolyn Tett and Tabea Schneider for useful discussions. Charlie Walsh, Rachael Kramp, David Clark, James Oberleitner, Surya Venkatesan, and Shivam Chand assisted in data collection.

