## supplementary material for "Pace of life predicts parasite resistance and fecundity tolerance, but not mortality tolerance, among Trinidadian guppies, *Poecilia reticulata*"

#### Contents


**1 Introduction** 1

**2 Import data and create new data frames** 1

**3 Checking the data** 2

**4 Question 1: Do the courses differ in resistance?** 3

**5 Question 2: Do the courses differ in fecundity tolerance?** 5

**6 Question 3: Do the courses differ in mortality tolerance?** 8

6.1 GLMM . . . . . . . . . . 8

6.2 Survival analysis . . . . . . . . . . 11


### 1 Introduction

We used the datasheet 'Inf_Dissections_Trini.csv' in this analysis, which contains the following variables:

**MaternalID:** Identify of the mother
**DateDissected:** Date sample was dissected in YYYYMMDD format **OrderDissected:** For each day of dissection, order dissection occurred in (1 = 1st sample dissected on that date, 2 = 2nd, etc.)
**OffspringNumber:** Number of offspring; since each offspring gets its own row, should be either 1 (with a stage in the next column) or 0 (mother had no offspring)
**OffspringDevelopmentalStage:** Development stage (A-J) of offspring
**offstage:** As above, but with letters converted to numbers - A=1, B=2.. etc. If no offspring were found, NA.
**offstage0:** As above, but with letters converted to numbers - A=1, B=2.. etc. If no offspring were found, 0.
**DissectorInitials:** Initials of dissectors (CW = Charlie Walsh, NT = Natalie Tepox)
**Batch:** Infections were conducted in batches 1-3
**Population:** Population of fish
**River:** River the progenitors of these fish were originally collected from **Course:** River course of collection **AUC:** Area under the curve of parasite over time
**D11AUC:** Area under the curve of parasite over time up to day 11 of infection
**InfDeathDays:** Number of days between infection and death (including euthanasia at end of experiment)
**PreDate:** Date in YYYYMMDD of weighing and lengthing fish before infection
**PreWeight:** Weight of fish in grams before infection
**PreLength:** Length of fish in centimeters before infection
**PreSMI:** Scaled mass length from before infection based on linear model predicting weight by length using preinfection metrics
**PostDate:** Date in YYYYMMDD of weighing and lengthing fish fater infection
**PostWeight:** Weight of fish in grams after infection
**PostLength:** Length of fish in centimeteres after infection
**PostSMI:** Scaled mass index from after infection based on linear model predicting weight by length using

preinfection metrics
**DeltaSMI:** PostSMI - PreSMI
**DoseDate:** Day fish was infected in YYYYMMDD
**Dose:** Total number of worms on fish for initial infection
**DeathDate:** Day of death in YYYYMMDD–for fish that lasted until the last day of the experiment, date of the last day and when they were euthanized
**PrematureDeath:** Y if fish died before end of eexperiment; N if fish was euathanized at end of experiment

#### 2 Import data and create new data frames

Our datasheet is organized so that each offspring has its own row. Here, we use the raw data to add a variable:

**totaloffspring:** adds all offspring attributed to that unique maternal ID to give a straightforward count of offspring.

We then create a new dataframe with only one row per mother.

```
DFIMP <- read.csv("Inf_Dissections_Trini.csv")
DFIMP$Maternal_ID <- as.factor(DFIMP$Maternal_ID)

ocount <- ddply(DFIMP, c("Maternal_ID"),summarise,
                   totaloffspring = sum(Offspring_Number))

dfall<-merge(ocount, DFIMP, by="Maternal_ID")
dfall$rD11AUC<-rescale(dfall$D11AUC)

#data with one observation
dfu <- dfall %>%
  arrange(Maternal_ID) %>%
distinct(Maternal_ID, .keep_all = TRUE)
```

#### 3 Checking the data

Looking at the correlations among variables to confirm our predictors are ok to include in the same model.

```
ggpairs(dfu,
             columns = c("AUC", "D11AUC", "InfDeathDays",
                         "totaloffspring", "offstage", "offstage0",
                         "Course", "PreSMI", "PreLength"))
```

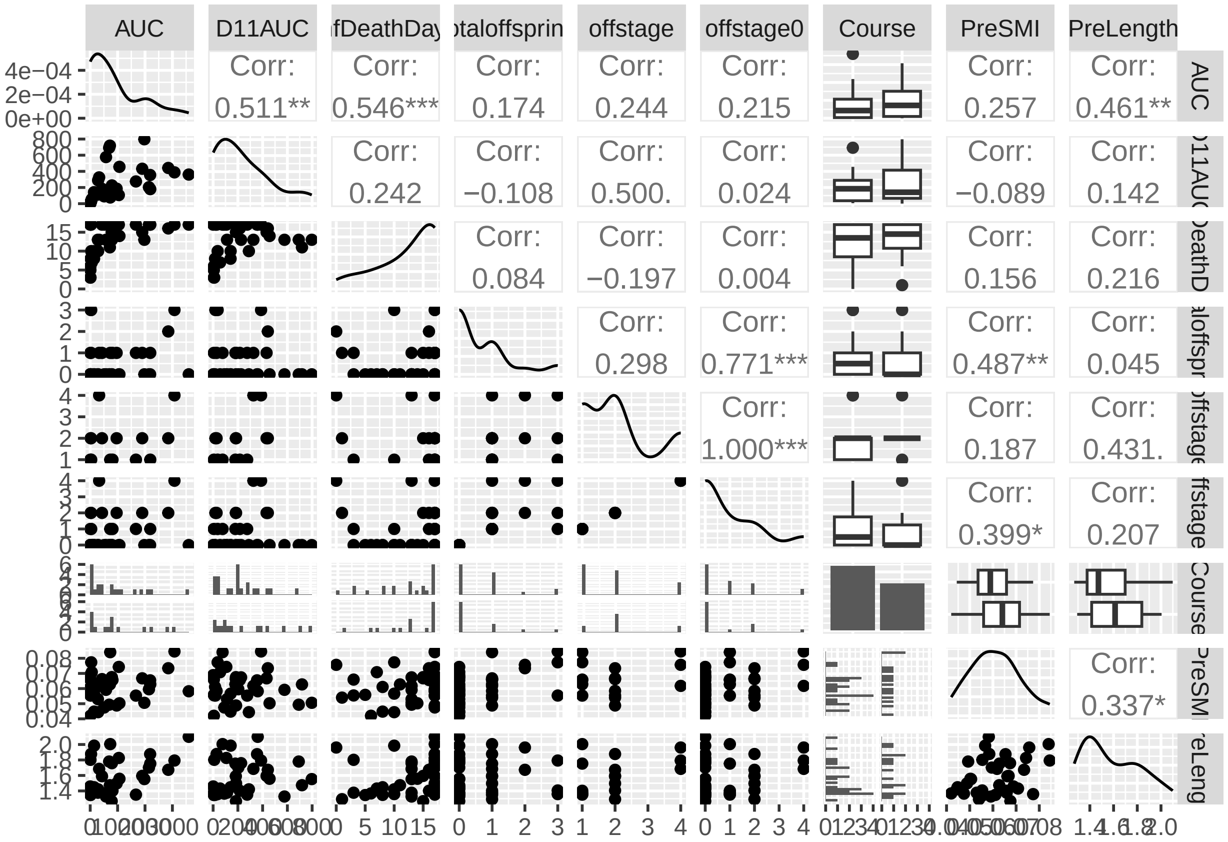


### 4 Question 1: Do the courses differ in resistance?

Lower courses invest less in resistance/get higher parasite loads.

```
names(dfu)
```

```
##  [1] "Maternal_ID"                "totaloffspring"
##  [3] "Date_Dissected"             "Order_Dissected"
##  [5] "Offspring_Number"           "Offspring_Development_Stage"
##  [7] "offstage"                   "offstage0"
##  [9] "Dissector_Initials"         "Batch"
## [11] "Population"                 "River"
## [13] "Course"                     "AUC"
## [15] "D11AUC"                     "InfDeathDays"
## [17] "PreDate"                    "PreWeight"
## [19] "PreLength"                  "PreSMI"
## [21] "PostDate"                   "PostWeight"
## [23] "PostLength"                 "PostLengthResid"
## [25] "PostSMI"                    "DeltaSMI"
## [27] "DoseDate"                   "Dose"
## [29] "DeathDate"                  "DonorID"
## [31] "PrematureDeath"             "rD11AUC"
```

```
#dfu2<-subset(dfu, "Maternal_ID"!="B1_UY_F4")# to remove the female
#that gave birth to 1 offspring during her infection

M1 <- lmer(log(D11AUC) ~ PreLength+
```

```r
                      totaloffspring+
                     Course+
                    InfDeathDays+
                    (1|DonorID)+
                    (1|River)+
                    (1|Population),
           data = dfu, na.action=na.omit)
```

```
## boundary (singular) fit: see help('isSingular')
```

```r
summary(M1)
```

```
## Linear mixed model fit by REML ['lmerMod']
## Formula: log(D11AUC) ~ PreLength + totaloffspring + Course + InfDeathDays +
##     (1 | DonorID) + (1 | River) + (1 | Population)
##    Data: dfu
##
## REML criterion at convergence: 123.4
##
## Scaled residuals:
##      Min       1Q   Median       3Q      Max
## -2.04532 -0.51052 -0.04227  0.59286  2.03910
##
## Random effects:
##  Groups     Name        Variance  Std.Dev.
##  DonorID    (Intercept) 1.297e+00 1.139e+00
##  Population (Intercept) 1.244e-09 3.527e-05
##  River      (Intercept) 1.126e+00 1.061e+00
##  Residual               1.267e+00 1.125e+00
## Number of obs: 36, groups:  DonorID, 6; Population, 5; River, 3
##
## Fixed effects:
##                Estimate Std. Error t value
## (Intercept)    -0.85415    1.87124  -0.456
## PreLength       2.79244    1.09607   2.548
## totaloffspring -0.10976    0.25315  -0.434
## CourseUpper    -1.72732    0.70613  -2.446
## InfDeathDays    0.15720    0.05347   2.940
##
## Correlation of Fixed Effects:
##             (Intr) PrLngt ttlffs CrsUpp
## PreLength   -0.836
## totlffsprng -0.151  0.159
## CourseUpper  0.054 -0.167  0.329
## InfDeathDys -0.100 -0.197 -0.388 -0.321
## optimizer (nloptwrap) convergence code: 0 (OK)
## boundary (singular) fit: see help('isSingular')
```

```r
Anova(M1)
```

```
## Analysis of Deviance Table (Type II Wald chisquare tests)
##
## Response: log(D11AUC)
##                 Chisq Df Pr(>Chisq)
## PreLength      6.4906  1   0.010844 *
```

```
## totaloffspring 0.1880  1   0.664592
## Course         5.9839  1   0.014437 *
## InfDeathDays   8.6443  1   0.003281 **
## ---
## Signif. codes:  0 '***' 0.001 '**' 0.01 '*' 0.05 '.' 0.1 ' ' 1
```

```r
simulateResiduals(M1, plot = T)
```

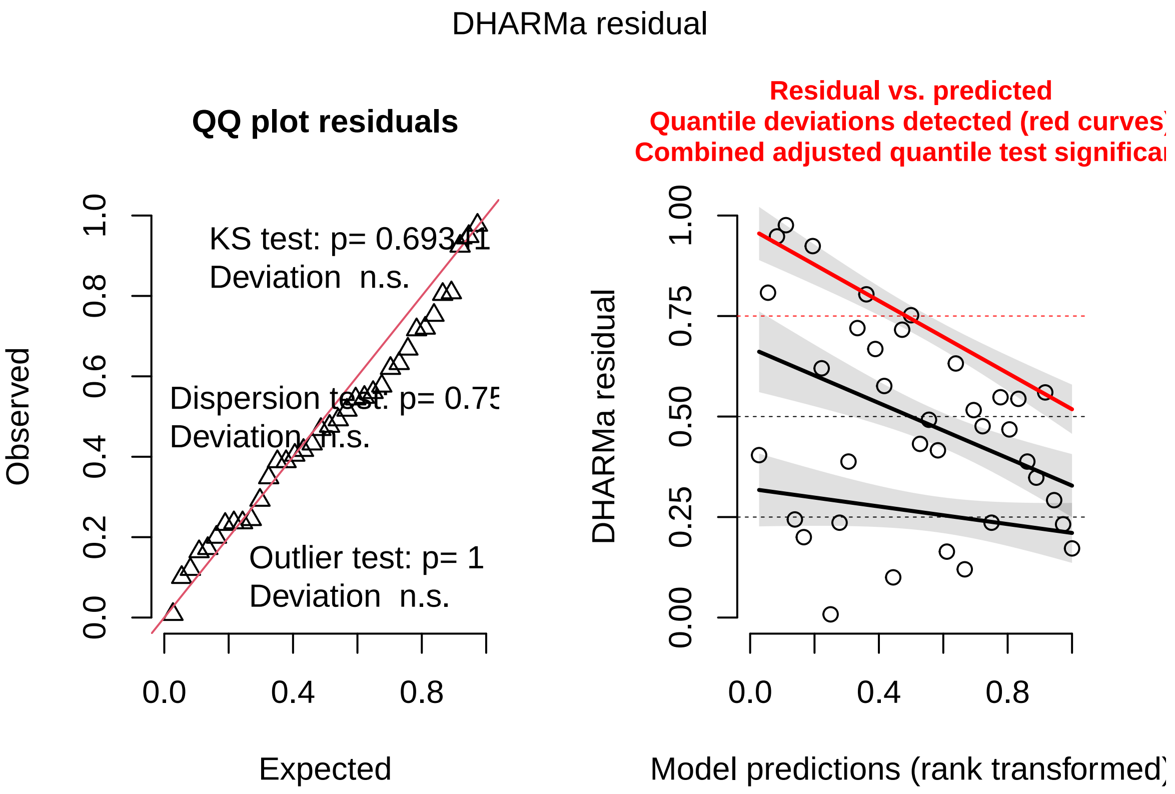


```
## Object of Class DHARMa with simulated residuals based on 250 simulations with refit = FALSE . See ?DI
##
## Scaled residual values: 0.432 0.668 0.236 0.348 0.56 0.388 0.172 0.292 0.416 0.804 0.404 0.72 0.548 (
```

```r
#variance calculation
#DonorID: 1.297
#River: 1.126
#Residual: 1.267
#Population: negligible

1.297/(1.297+1.126+1.267)
```

```
## [1] 0.3514905
```

```r
1.126/(1.297+1.126+1.267)
```

```
## [1] 0.3051491
```

### 5 Question 2: Do the courses differ in fecundity tolerance?

For this analysis - it's at the level of the individual mother - we're testing essentially "are there maternal attributes such as her population of origin, her infection severity, her size.. etc that predict the number of offspring we found in her?"

```r
M2 <- glmer(totaloffspring ~ PreLength +
                     Course*D11AUC+
                    InfDeathDays+
              (1|DonorID)+
              (1|Population)+
              (1|River),
                   family=poisson(link="log"), data = dfu, na.action=na.omit)
summary(M2)
```

```
## Generalized linear mixed model fit by maximum likelihood (Laplace
##   Approximation) [glmerMod]
##  Family: poisson  ( log )
## Formula: totaloffspring ~ PreLength + Course * D11AUC + InfDeathDays +
##     (1 | DonorID) + (1 | Population) + (1 | River)
##    Data: dfu
##
##      AIC      BIC   logLik deviance df.resid
##     82.2     96.5    -32.1     64.2       27
##
## Scaled residuals:
##     Min      1Q  Median      3Q     Max
## -1.0485 -0.6405 -0.3637  0.3855  2.6665
##
## Random effects:
##  Groups     Name        Variance Std.Dev.
##  DonorID    (Intercept) 0        0
##  Population (Intercept) 0        0
##  River      (Intercept) 0        0
## Number of obs: 36, groups:  DonorID, 6; Population, 5; River, 3
##
## Fixed effects:
##                      Estimate Std. Error z value Pr(>|z|)
## (Intercept)         -2.806831   1.924237  -1.459  0.14466
## PreLength            0.691594   1.264918   0.547  0.58455
## CourseUpper         -2.135769   0.935413  -2.283  0.02242 *
## D11AUC              -0.006411   0.002764  -2.320  0.02035 *
## InfDeathDays         0.178181   0.066695   2.672  0.00755 **
## CourseUpper:D11AUC  0.007696   0.003389   2.271  0.02314 *
## ---
## Signif. codes:  0 '***' 0.001 '**' 0.01 '*' 0.05 '.' 0.1 ' ' 1
##
## Correlation of Fixed Effects:
##             (Intr) PrLngt CrsUpp D11AUC InfDtD
## PreLength   -0.869
## CourseUpper  0.488 -0.536
## D11AUC       0.310 -0.374  0.608
## InfDeathDys -0.353 -0.099 -0.231 -0.276
## CrsU:D11AUC -0.383  0.412 -0.806 -0.884  0.299
## optimizer (Nelder_Mead) convergence code: 0 (OK)
## boundary (singular) fit: see help('isSingular')
```

```r
Anova(M2)
```

```
## Analysis of Deviance Table (Type II Wald chisquare tests)
```

```
##
## Response: totaloffspring
##                 Chisq Df Pr(>Chisq)
## PreLength      0.2989  1   0.584551
## Course         0.5847  1   0.444458
## D11AUC         0.4463  1   0.504081
## InfDeathDays   7.1375  1   0.007549 **
## Course:D11AUC  5.1578  1   0.023142 *
## ---
## Signif. codes:  0 '***' 0.001 '**' 0.01 '*' 0.05 '.' 0.1 ' ' 1
```

```r
simulateResiduals(M2, plot = T)
```

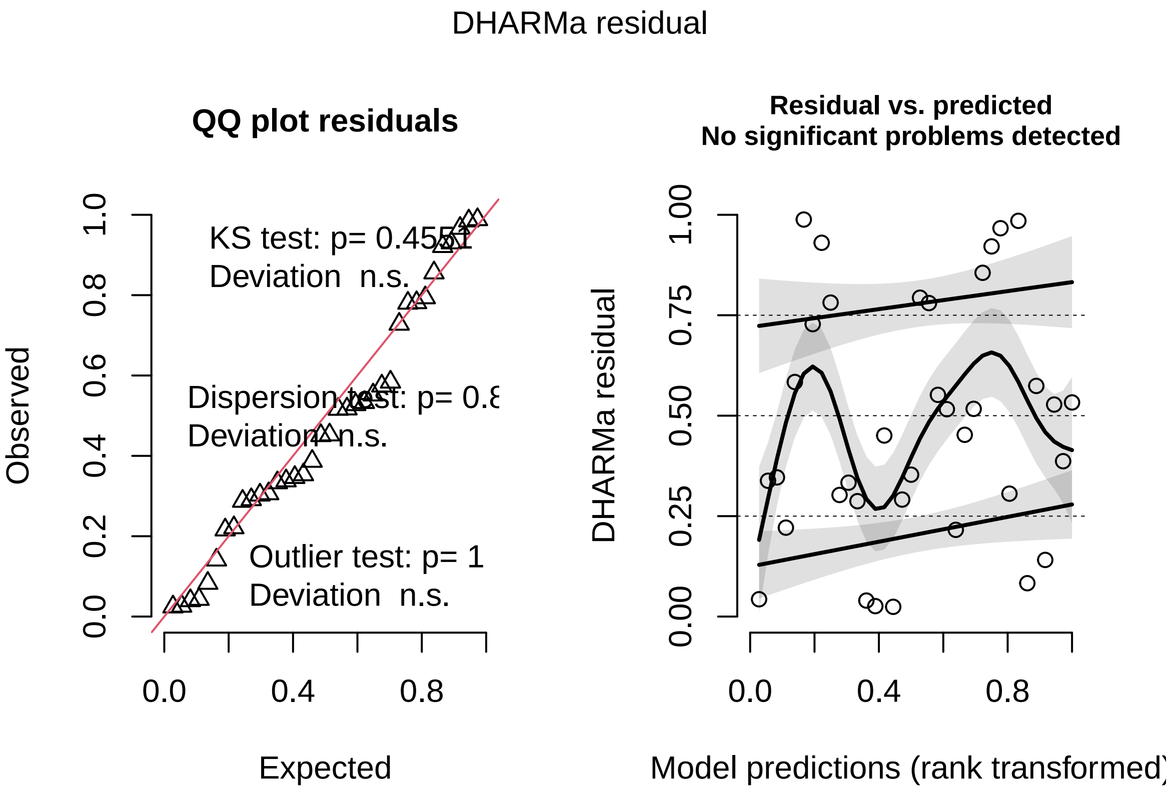


```
## Object of Class DHARMa with simulated residuals based on 250 simulations with refit = FALSE . See ?DI
##
## Scaled residual values: 0.3059428 0.3026633 0.5518062 0.5277377 0.04307911 0.5740832 0.3531546 0.140
```

```r
#post hoc test
M2.post <- test(emtrends(M2, specs = "Course", var = "D11AUC"))
summary(M2.post)
```

```
##  Course D11AUC.trend      SE  df z.ratio p.value
##  Lower      -0.00641 0.00276 Inf  -2.320  0.0203
##  Upper       0.00128 0.00160 Inf   0.802  0.4224
```

### 6 Question 3: Do the courses differ in mortality tolerance?

#### 6.1 GLMM

Overall no - there is no difference in probability of dying, and no difference in how infection loads contribute to this probability.

```
#Creating new variable of whether or not they died prematurely
dfu$PD <- ifelse(dfu$PrematureDeath == "Y", 1, 0)  #

M3<-glmer(PD~
              totaloffspring+
              Course+
              D11AUC+
              PreLength+
              D11AUC:Course+
            (1|Population)+
              (1|River),
            data=dfu,
            family=binomial(link="logit"))

summary(M3)
```

```
## Generalized linear mixed model fit by maximum likelihood (Laplace
##   Approximation) [glmerMod]
##  Family: binomial  ( logit )
## Formula: PD ~ totaloffspring + Course + D11AUC + PreLength + D11AUC:Course +
##     (1 | Population) + (1 | River)
##    Data: dfu
##
##      AIC      BIC   logLik deviance df.resid
##     41.4     54.0    -12.7     25.4       28
##
## Scaled residuals:
##      Min       1Q   Median       3Q      Max
## -2.85904  0.06984  0.24655  0.31793  1.79109
##
## Random effects:
##  Groups     Name        Variance  Std.Dev.
##  Population (Intercept) 4.708e-17 6.862e-09
##  River      (Intercept) 3.190e-17 5.648e-09
## Number of obs: 36, groups:  Population, 5; River, 3
##
## Fixed effects:
##                       Estimate Std. Error z value Pr(>|z|)
## (Intercept)          12.929047   4.693671   2.755  0.00588 **
## totaloffspring        0.013745   0.631729   0.022  0.98264
## CourseUpper           0.289717   1.776322   0.163  0.87044
## D11AUC                0.002544   0.003773   0.674  0.50007
## PreLength            -7.557644   2.895887  -2.610  0.00906 **
## CourseUpper:D11AUC    0.001820   0.006663   0.273  0.78478
## ---
## Signif. codes:  0 '***' 0.001 '**' 0.01 '*' 0.05 '.' 0.1 ' ' 1
##
## Correlation of Fixed Effects:
```

```
##              (Intr) ttlffs CrsUpp D11AUC PrLngt
## totlffsprng -0.221
## CourseUpper  0.172  0.202
## D11AUC       0.059  0.135  0.567
## PreLength   -0.968  0.114 -0.345 -0.252
## CrsU:D11AUC  0.056 -0.302 -0.762 -0.586  0.076
## optimizer (Nelder_Mead) convergence code: 0 (OK)
## boundary (singular) fit: see help('isSingular')
```

```r
Anova(M3)
```

```
## Analysis of Deviance Table (Type II Wald chisquare tests)
##
## Response: PD
##                  Chisq Df Pr(>Chisq)
## totaloffspring 0.0005  1    0.98264
## Course         0.3280  1    0.56683
## D11AUC         1.0594  1    0.30334
## PreLength      6.8110  1    0.00906 **
## Course:D11AUC  0.0746  1    0.78478
## ---
## Signif. codes:  0 '***' 0.001 '**' 0.01 '*' 0.05 '.' 0.1 ' ' 1
```

```r
simulateResiduals(M3, plot = T)
```

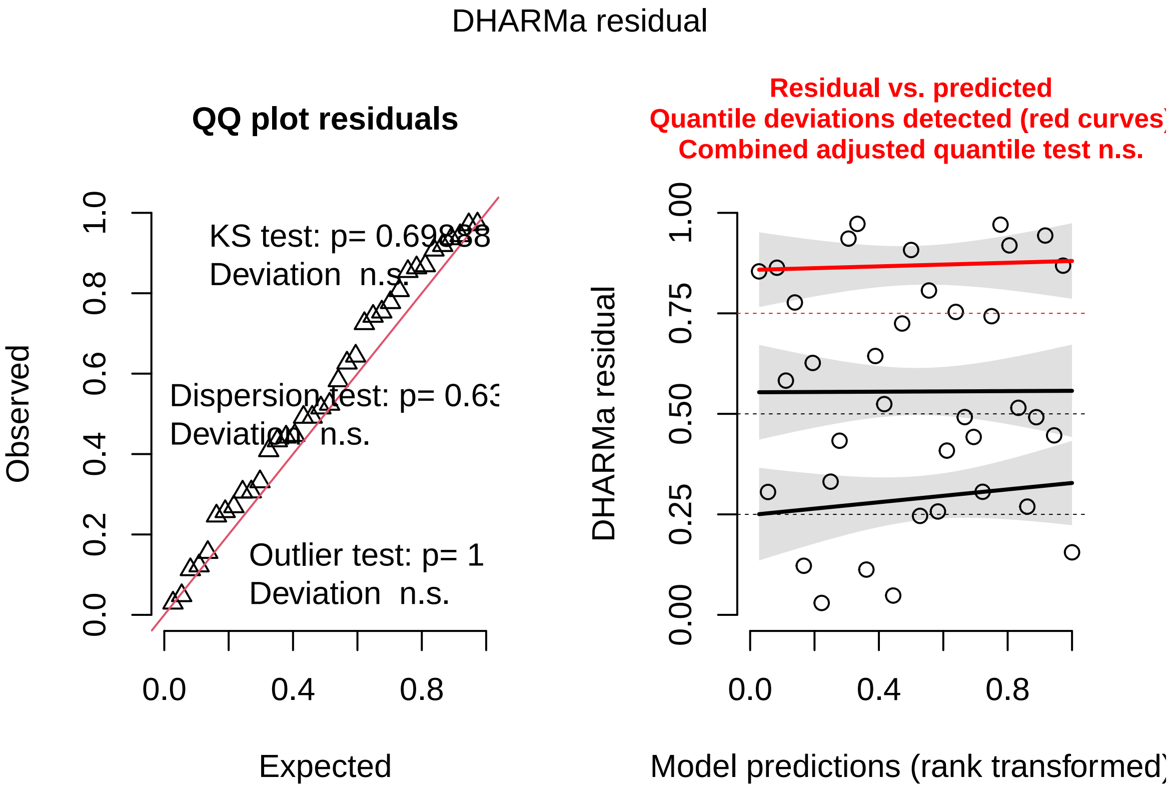


```
## Object of Class DHARMa with simulated residuals based on 250 simulations with refit = FALSE . See ?DH
##
## Scaled residual values: 0.7428972 0.2462121 0.4425258 0.1126584 0.9725422 0.0480893 0.8543842 0.0296
```

```r
# more likely to die if smaller
visreg(M3, "PreLength", scale = "response")
```

```
## Conditions used in construction of plot
## totaloffspring: 0
## Course: Lower
## D11AUC: 184
## Population: Lower Guanpo
## River: Aripo
```

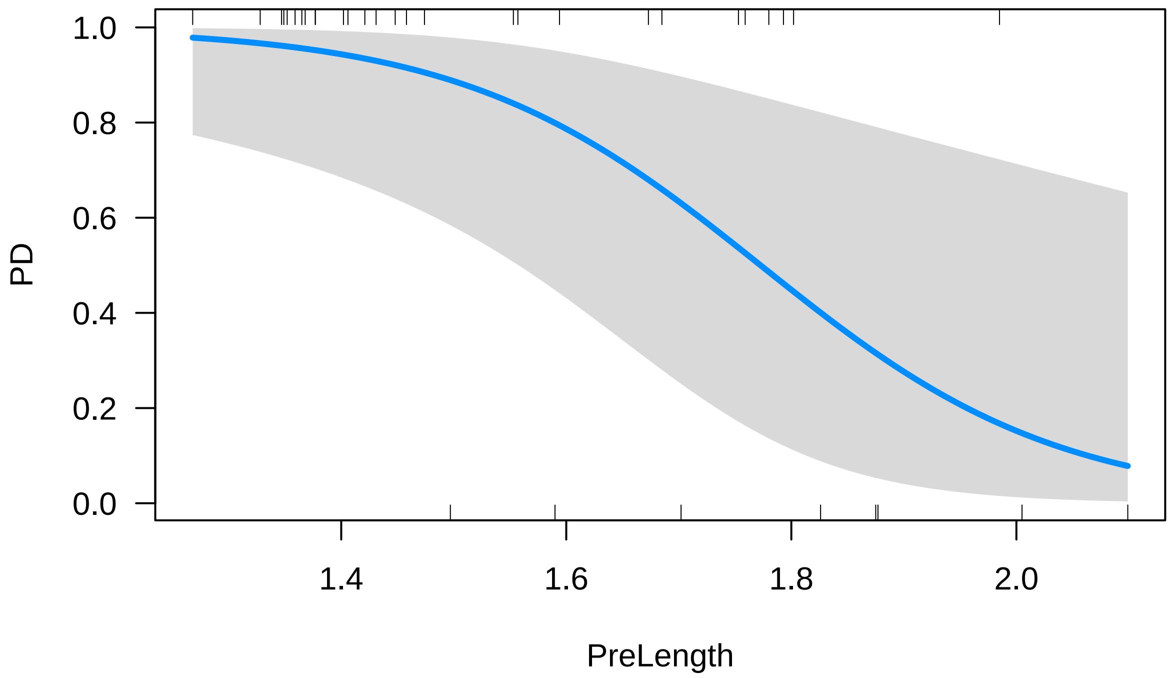


```
visreg(M3, "D11AUC", "Course", scale = "response")
```

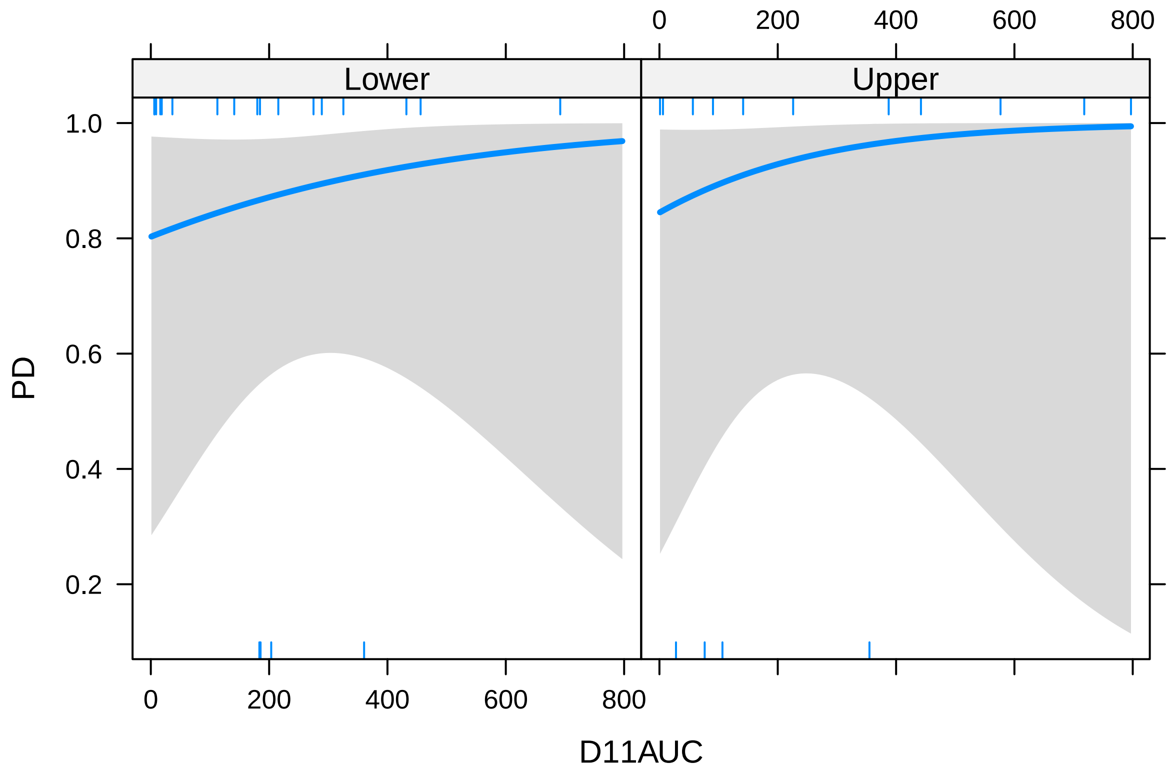

#### 6.2 Survival analysis

```
#Survival analysis just in case we need it

library(survival)
library(survminer)
library(coxme)
library(ade4)

#Analysis type Kaplan/Meier o Cox Model

dfu$surv_time <- dfu$InfDeathDays
dfu$status <- ifelse(dfu$InfDeathDays < 17, 1, 0)  # 1 = murio antes del dia 17, 0 = censurada (viva ha

cox_dfio <- coxph(Surv(surv_time, status) ~ Course*D11AUC, data = dfu)
summary(cox_dfio)
```

```
## Call:
## coxph(formula = Surv(surv_time, status) ~ Course * D11AUC, data = dfu)
##
##   n= 36, number of events= 22
##    (2 observations deleted due to missingness)
##
##                          coef exp(coef)  se(coef)      z Pr(>|z|)
## CourseUpper        -1.304855  0.271212  0.788611 -1.655    0.098 .
## D11AUC             -0.001685  0.998316  0.001991 -0.846    0.397
## CourseUpper:D11AUC  0.003585  1.003591  0.002411  1.486    0.137
## ---
## Signif. codes:  0 '***' 0.001 '**' 0.01 '*' 0.05 '.' 0.1 ' ' 1
##
##                    exp(coef) exp(-coef) lower .95 upper .95
## CourseUpper           0.2712     3.6872   0.05782     1.272
## D11AUC                0.9983     1.0017   0.99443     1.002
## CourseUpper:D11AUC    1.0036     0.9964   0.99886     1.008
##
## Concordance= 0.628  (se = 0.079 )
## Likelihood ratio test= 3.22  on 3 df,   p=0.4
## Wald test            = 2.98  on 3 df,   p=0.4
## Score (logrank) test = 3.08  on 3 df,   p=0.4
```

```
Anova(cox_dfio)
```

```
## Analysis of Deviance Table (Type II tests)
##               LR Chisq Df Pr(>Chisq)
## Course         0.63701  1     0.4248
## D11AUC         0.27140  1     0.6024
## Course:D11AUC  2.35316  1     0.1250
```

```
ggsurvplot(survfit(Surv(surv_time, status) ~ Course, data = dfu),
           data = dfu,
           risk.table = TRUE,
           pval = TRUE,
           xlab = "Post Infection Days",
           ylab = "Survival Probability",
           palette = "Dark2")
```

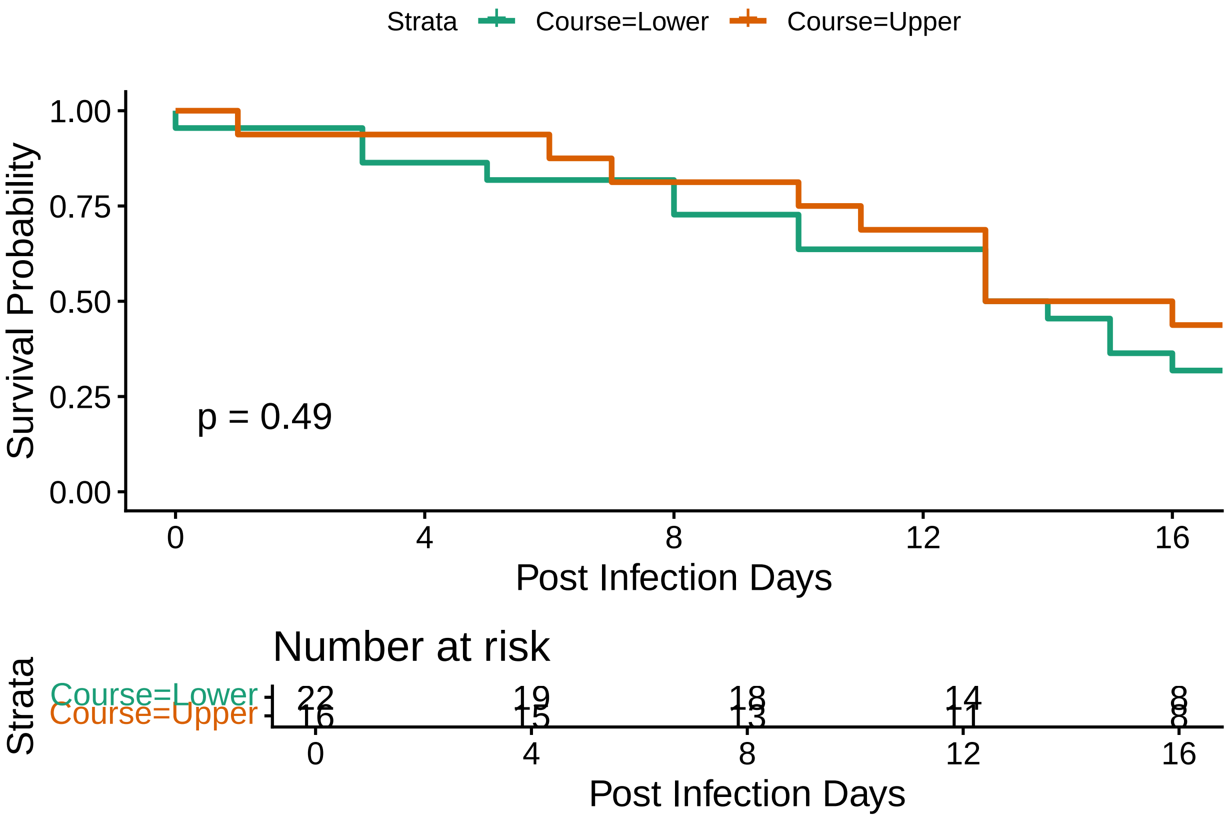


```
# Can use a mixed effects cox model with the same predictors as the binomial mortality glm
coxme_dfio <- coxme(Surv(surv_time, PD) ~ totaloffspring +
                      Course +
                      D11AUC +
                      PreLength +
                      D11AUC:Course +
                      (1|Population) + (1|River), data = dfu)
Anova(coxme_dfio) # same results
```

```
## Analysis of Deviance Table (Type II tests)
##
## Response: Surv(surv_time, PD)
##                Df  Chisq Pr(>Chisq)
## totaloffspring  1 0.7988    0.37145
## Course          1 0.1102    0.73992
## D11AUC          1 0.6683    0.41365
## PreLength       1 5.3966    0.02018 *
## Course:D11AUC   1 0.8203    0.36510
## ---
## Signif. codes:  0 '***' 0.001 '**' 0.01 '*' 0.05 '.' 0.1 ' ' 1
```
